# Macrophages drive a fibrogenic gene program of periductal fibroblasts in pediatric primary sclerosing cholangitis

**DOI:** 10.1101/2025.08.14.670195

**Authors:** Yunguan Wang, David Adeleke, Xiangfei Xie, Zi Yang, Annika Yang von Hofe, Manavi Singh, Astha Malik, Ramesh Kudira, Cyd Castro-Rojas, Liva Pfuhler, Mosab Alquraish, Pamela Sylvestre, Jonathan R. Dillman, Andrew T. Trout, Emily Miraldi, Alexander G. Miethke

## Abstract

Primary sclerosing cholangitis (PSC) is an autoimmune, cholestatic liver disease characterized by inflammation and fibrosis surrounding bile ducts. The cellular crosstalk driving periductal fibrosis remains poorly defined. This study applied a multi-omics approach integrating MERSCOPE spatial transcriptomics, bulk RNA-seq, and SomaScan proteomics to characterize fibrotic periductal regions and their cell–cell communications. Macrophages (MP) expressing moderate-to-high CD163 were found co-localized with cholangiocytes, T cells, and collagen-producing hepatic stellate cells (HSC). Cell niche analysis identified periductal regions with elevated fibrotic signals, in which cell–cell communication analysis revealed MP–HSC interactions involving 17 fibrotic driver genes in MP (e.g., IFNGR1, CSF1R, CD163) and six fibrotic effector genes in HSC (e.g., COL1A2, VCAN, MMP2). In validation analyses, bulk RNA-seq data showed higher driver and effector gene scores in PSC with established fibrosis compared to early-stage PSC and autoimmune hepatitis (AIH). Plasma proteins encoded by MP driver genes were elevated in autoimmune liver disease (AILD) and in patients with elevated (≥3.29 kPa) liver stiffness on MR elastography. These findings suggest that macrophages engage in localized crosstalk with HSC, activating fibrotic gene programs and contributing to periductal fibrosis in PSC, thereby identifying potential molecular targets for therapeutic intervention.

## Introduction

Primary sclerosing cholangitis (PSC) is a chronic, idiopathic cholestatic liver disease characterized by inflammation and fibrosis of the bile ducts. Although the exact etiology of PSC remains unclear, genome-wide association studies (GWAS) have implicated genes highly expressed in lymphocytes and myeloid cells—such as HLA genes, FCRL3, INAVA, PRDM1, CCR6, CD226, and IL12RB1—as risk factors for developing PSC (1). PSC is a slowly, but relentlessly progressive disease in children and adolescents with 38% and 25% developing portal hypertensive and biliary complications, respectively, within 10 years of disease onset and 14% of patients requiring liver transplantation to prolong survival, as shown in a large, multicenter, retrospective cohort study (2). Unique to children and adolescents is the overlap with autoimmune hepatitis (AIH) in 33% of patients with PSC, whereas patients of all ages are frequently diagnosed with concomitant inflammatory bowel disease (IBD). Current treatments focus on managing symptoms and complications, but no approved therapy effectively halts hepatobiliary fibrosis progression (3).

The development of hepatobiliary fibrosis involves complex interactions between several cell types, including cholangiocytes, fibroblasts (portal fibroblasts, PF, and hepatic stellate cells, HSC), and immune cells. In mouse models of sclerosing cholangitis, injury to bile duct epithelial cells activate nearby fibroblasts which then migrate to bile ducts accelerating liver fibrosis progression (4). Both PF and recruited HSC contribute to biliary fibrosis by adopting a myofibroblast phenotype characterized by ACTA2 upregulation and excessive collagen production in mouse models of PSC (5). Macrophages also play a critical role in PSC pathogenesis. These cells, known for their diverse functions, have both pro-inflammatory and fibrogenic properties (6). Macrophages promote inflammation and liver damage by secreting cytokines that recruit and activate other immune cells. Additionally, macrophages contribute directly to fibrosis by activating HSC and producing fibrogenic cytokines such as TGF-β (7). While macrophages have been identified as key drivers of hepatobiliary fibrosis in PSC, their precise roles across macrophage subpopulations remain poorly understood. Gores et al. demonstrated that inhibiting macrophage recruitment reduced peribiliary fibroinflammation in an Mdr2-/-mouse model of sclerosing cholangitis (7). In contrast, in a cholestatic injury model involving bile duct ligation and dietary 3,5-diethoxycarbonyl-1,4-dihydrocollidine (DDC), macrophage depletion during early cholestasis induction had no significant impact on fibrosis progression (8, 9).

Recent studies investigating PSC liver immunology in humans revealed macrophages exist in distinct subtypes, including Kupffer cells (KC), monocyte-like macrophages (MoMP), and lipid-associated macrophages (LAM-like) (9, 10). Spatially resolved transcriptomics (SRT) analysis using the Visium platform showed that these macrophage subtypes are distributed differently across the portal-central axis and between scarring and non-scarring regions. However, the limited resolution of the Visium platform leaves gaps in our understanding of how macrophages are spatially organized within fibrotic niches and how their transcriptomic heterogeneity contributes to local fibrotic gene programs.

To address these gaps, we employed a multi-omics approach integrating single-cell spatially resolved transcriptomics (scSRT), RNA-seq, and plasma proteomics data from AILD patients and healthy donors (n = 2, 64, and 122, respectively, across the differ platforms) to investigate the role of macrophages in fibroblast activation (Fig. 1). Using unsupervised cell niche analysis, we identified fibrotic regions in liver biopsies and characterized the macrophage populations enriched within them. We then prioritized key cell-cell communication (CCC) genes that may mediate interactions between macrophages and fibrogenic fibroblasts. We hypothesized that gene programs identified through CCC in scSRT data would also exhibit strong associations in non-spatial data. Consistent with this, we confirmed that fibrogenic gene expression in fibroblasts correlates with CCC-associated driver genes in macrophages in PSC scRNA-seq data and that these fibrotic gene signatures are linked to advanced fibrosis stages in pediatric PSC.

**Figure 1.**
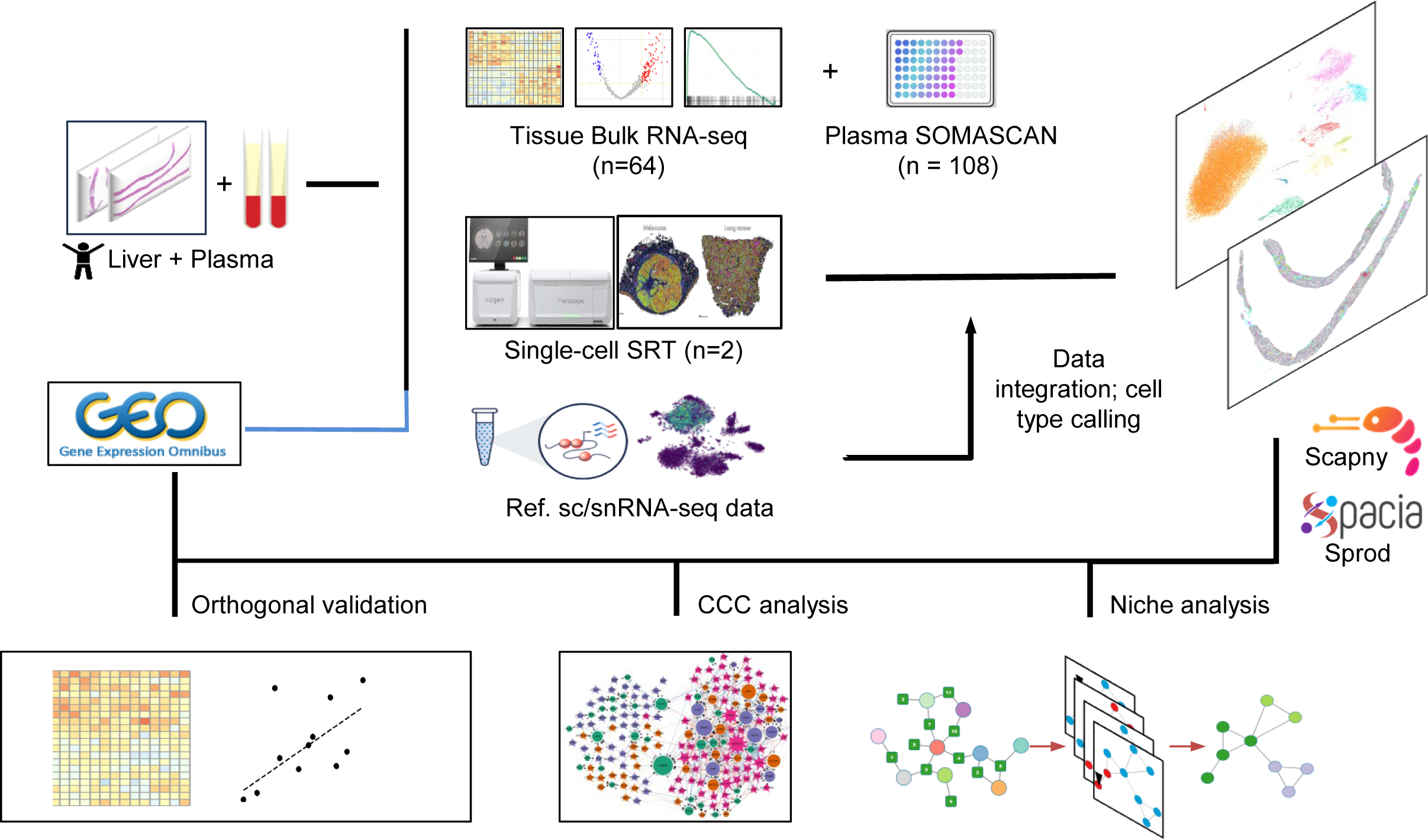
Analytical workflow for scSRT data analysis, integration, CCC prediction, and orthogonal validation.

## Results

### Macrophage activity is correlated with advanced fibrosis in AILD

To investigate the association between macrophage activation and liver fibrosis, we performed RNA sequencing of cryopreserved tissue samples from 64 clinically indicated liver biopsies of pediatric patients with PSC or AIH. These samples were collected at the Center for Autoimmune Liver Disease (CALD) at Cincinnati Children’s Hospital Medical Center and clinical data are summarized in **Table 1**. Given the potential biases introduced by tissue heterogeneity and bulk RNA-seq data, we applied unsupervised clustering to identify common proinflammatory and fibrogenic gene expression patterns. This analysis revealed three distinct clusters (**Fig. 2a**): cluster c1 was predominantly composed of PSC samples (68% PSC), compared to the other two clusters (Odds ratio =2.72, P-value = 0.10, **Fig. 2b**). Notably, cluster c1 was significantly enriched in patients with advanced fibrosis (Odds ratio = 5.000, p-value = 0.007), as defined by a METAVIR score ≥ 2 or an ISHAK score ≥ 3 (**Fig. 2b**). Patients in c1 exhibited more bile duct injuries, as evidenced by higher alkaline phosphatase (p = 0.002), total bilirubin (p < 0.001), and serum gamma-glutamyl transferase levels (GGT) (p = 0.08) (**Fig. 2c**). To further explore the relationship between gene expression and fibrosis, we performed Gene Set Enrichment Analysis (GSEA) using genes differentially expressed in cluster c1. We found upregulation of pathways involved in TGF-beta signaling and collagen formation, while genes related to bile and lipid metabolism were among the most downregulated in patients from c1 compared with patients in c0 or c2 (**Fig. 2d**). Importantly, genes involved in macrophage activation, including *PLAUR*, *CXCL3*, *THBS1, AREG, NAMPT, IL1B, CD83 and CCL3*, were also upregulated in cluster c1 (**Fig. 2d**). To further assess macrophage activity in these AILD samples, we calculated the average expression of macrophage-related genes identified in a reference single-cell RNA sequencing dataset of PSC patients (9). Genes associated with macrophage activation—including those highly expressed in LAM and MoMP subpopulations—were significantly upregulated in the PSC-dominant cluster c1 (p-value < 0.001 for all comparisons, **Fig. 2e**). Genes associated with abundance of LAM included *SPP1, FABP5, CSTB, CD9, GPNMB* whereas *VCAN, S100A6, S100A4, LYZ, S100A9, S100A12 and S100A8* identified MoMP. Collectively, these results suggest that macrophage activity is elevated in patients with AILD and advanced fibrosis, potentially contributing to the activation of fibrotic gene programs.

**Figure 2.**
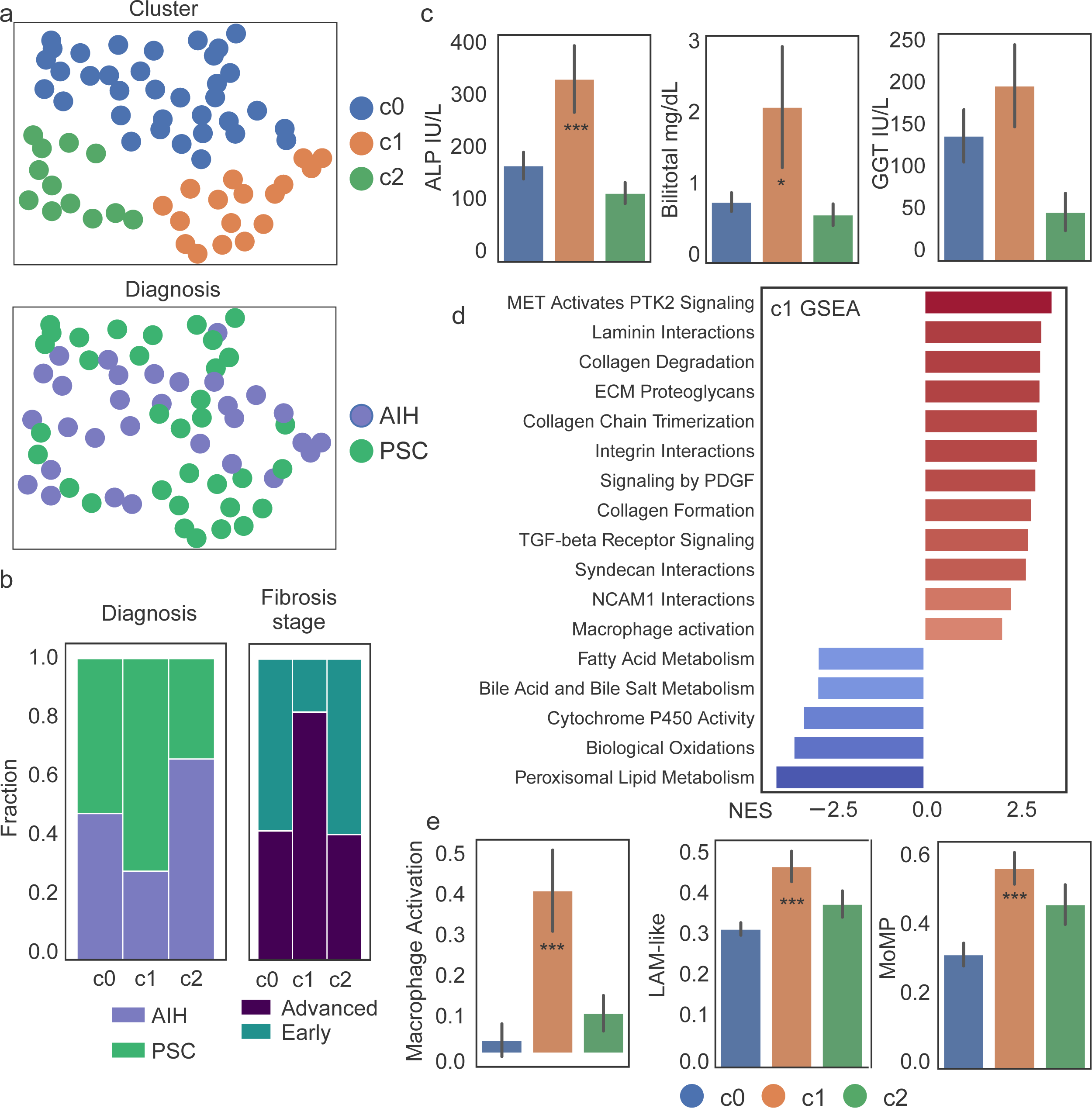
Evaluation of the association between macrophage gene programs and liver fibrosis in PSC. (a) Unsupervised clustering 64 AILD patient samples shown in embeddings calculated using the Uniform manifold approximation and projection (UMAP) algorithm. Samples were colored based on clusters (upper) or patient diagnosis at the time of data collection (lower). (b) Compositions of each patient sample cluster, based on diagnosis (left) or fibrosis stage (right). (c) Serum GGT (left), ALP (middle) and Bilirubin (right) levels in each patient cluster. (d) GSEA results calculated from the ranked differentially expressed genes in cluster 1 compared with c0 and c2. Pathways shown were selected based on an FDR q-value cutoff of 0.25, and colored based on the normalized enrichment scores (NES). Both the q-value and the NES were calculated using the gseapy Python package. (e) Average expression of genes involved in macrophage (MP) activation (left), and of marker genes for LAM, and MoMP (right) in each patient cluster. Values were normalized to a numeric range of 0-1. (d and e) Significance was obtained from the student’s t test comparing c1 vs the other patient clusters: ∗p <0.05, ∗∗p <0.01, ∗∗∗p <0.001. Standard errors were shown as error bars.

### Cell types identified by scSRT recapitulate the cellular landscape of PSC

To investigate the transcriptomic and spatial heterogeneity of these macrophage populations as putative drivers of fibrosis in PSC, we performed single-cell spatial transcriptomics (scSRT) using the VizGen MERSCOPE platform and a custom 400-gene liver immunology panel (Table S1). Two fresh-cut archived formalin-fixed paraffin-embedded (FFPE) slides from diagnostic liver biopsies of two patients with pediatric onset PSC with areas of focal periductal and bridging fibrosis were analyzed. At the time of the liver biopsy, participant A was a 10-year-old female with new diagnosis of PSC and possible overlap syndrome with AIH and past medical history of autoimmune pancreatitis and IBD. Participant B, 12-year-old male with a history of elevated liver function tests (LFTs) and abnormal liver imaging, was diagnosed with PSC based on the liver biopsy results (**Table 2**, **Fig. 3a and b**). Quality control of the scSRT data included a series of filtering steps based on metrics such as cell size, read counts, and doublet probability. A total of 93,879 cells were retained, with a median per-cell transcript count of 95. To mitigate dropout effects and improve segmentation, we incorporated data denoising and transcript-based re-segmentation in the preprocessing workflow, using Sprod (*1*) and Baysor (11), respectively.

**Figure 3.**
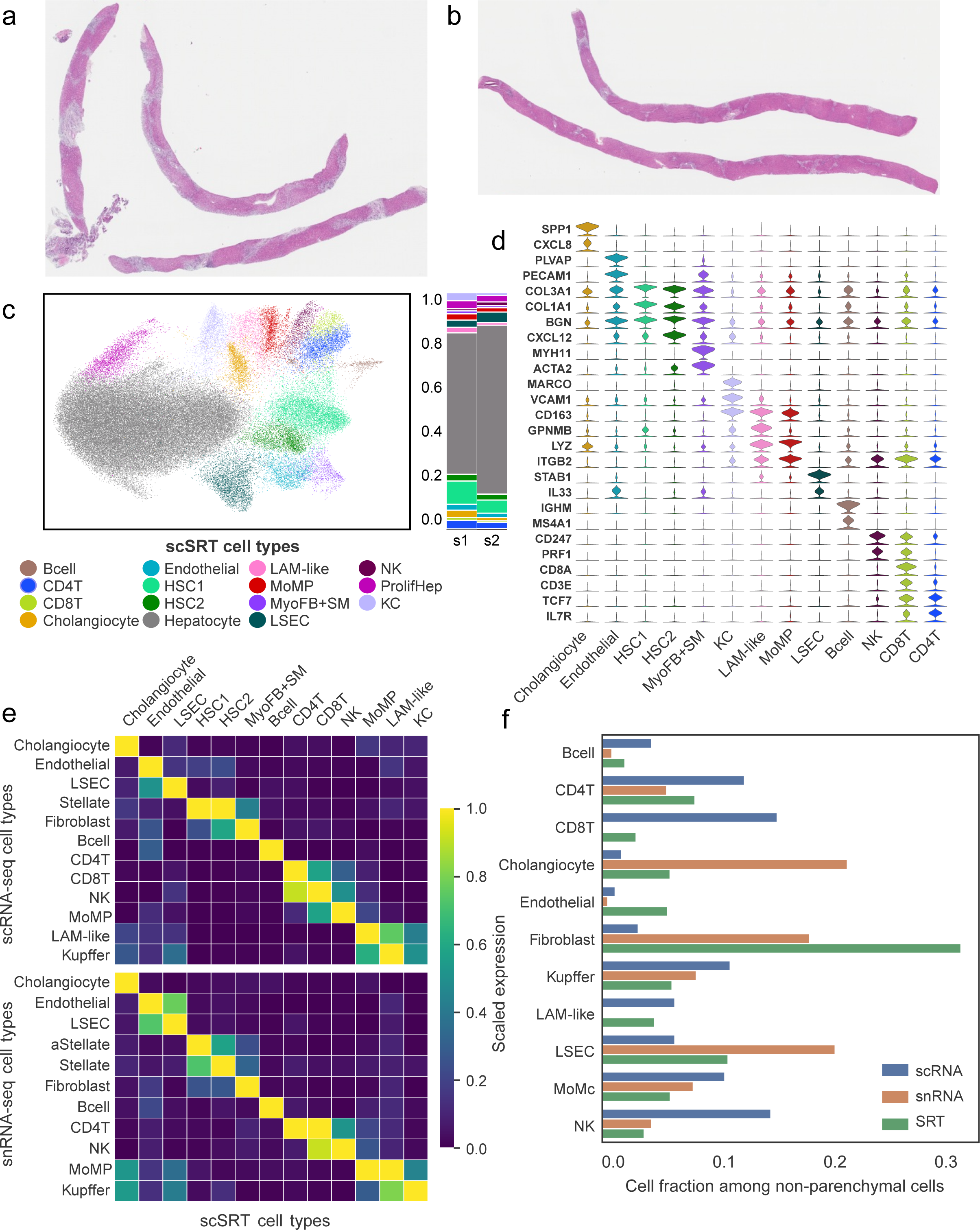
Evaluation of the cell types identified in scSRT PSC samples. (a and b) Overview of the two PSC samples used in this study. FFPE slides were stained with hematoxylin and eosin. Scale bar = 1mm. (c) PSC cell types identified using unsupervised clustering were shown in embeddings calculated using the UMAP algorithm. Samples were colored based on cell types. (d) Violin plot showing expression of selected marker genes of each cell type. Each violin was colored based on cell types shown in panel c. (d) Average expression of top 5 DEG from each scSRT cell type (rows) in scRNA-seq cell types (columns) in reference scRNA-seq (top) and snRNA-seq (bottom) datasets. Log1p(CPM) gene expression values were scaled to a range of 0 to 1. (f) Fractions of each cell type among all non-hepatocytes in scSRT and scRNA-seq data.

Next, we applied unsupervised clustering and identified major PSC-associated cell types using reference PSC cell type marker genes. These included hepatocytes, cholangiocytes, liver sinusoidal endothelial cells (LSEC), macrophages, T cells, and hepatic stellate cells (HSCs) (**Fig. 3c**). Notably, two HSC subpopulations emerged: HSC1, characterized by high expression of classical fibrosis genes *COL3A1* and *COL1A1*, and HSC2, which predominantly expressed *COL6A1* and *CXCL12*. Similarly, three macrophage populations were identified: LAM, MoMP, and KC (**Fig. 3d**). Interestingly, all three macrophage subsets exhibited moderate to high expression of the noninflammatory macrophage marker *CD163*. CD4 T cells highly expressed naïve/central-memory markers including *TCF7 and IL7R,* and tissue resident markers including *CD69* and *CXCR4.* This phenotype was consistent with the naïve-like T cells previously reported in PSC (12). CD8 T cells exhibited cytotoxic and central-memory phenotypes, with high expression of GZMK, PRF1, *TCF7* and *CD69* (**Fig. S1**). Additional differentially expressed genes (one vs rest) were summarized and reported in Supplementary Table 2.

To assess the concordance between cell types identified in scSRT and those from single-cell RNA sequencing (scRNA-seq), we reanalyzed scRNA-seq and single-nuclei (sn) RNA-seq data from 16 healthy subjects and patients with PSC or primary biliary cholangitis (PBC). We computed the UMAP projection of single cells from eight PSC patients, including seven with decompensated cirrhosis (DC), using only the 400 MERSCOPE genes and found that cell types remained well-separated (**Fig. S2**). Next, we evaluated the expression of marker genes from our scSRT cell types in the two reference datasets. As shown in **Fig. 3e**, marker genes identified from scSRT cell types including CD4/8 T cells, B cells, macrophages, hepatocytes and cholangiocytes were selectively expressed in the corresponding cell types from the scRNA-seq/snRNA-seq datasets. Based on the scRNAseq data set, HSC1 from the scSRT corresponded to activated stellate cells, while HSC2 expressed genes found in quiescent stellate cells. These results supported the robustness of the cell type annotations. We then compared the composition of cell types, excluding hepatocytes, between the two liver transcriptomic datasets from PSC patients. To facilitate interpretation, we grouped HSCs, myofibroblasts into a general fibroblast population. The sc/sn RNA-seq data contained a higher proportion of lymphoid and myeloid cells, whereas scSRT were enriched with cholangiocytes, endothelial cells, and fibroblasts. Notably, fibroblasts were ten times more abundant in the scSRT dataset compared to scRNA-seq, and nearly twice as abundant as in snRNA-seq, (**Fig. 3f**), despite the sc/sn RNA-seq dataset being primarily derived from highly fibrotic livers. This discrepancy in cell abundance highlights a key advantage of slide-based scSRT over droplet-based scRNA-seq platforms in capturing a more accurate cellular landscape, particularly for irregularly shaped cells that may be lost during scRNA-seq sample preparation (13, 14).

### Macrophage and fibroblast populations occupy distinct spatial regions

To investigate the transcriptomic and spatial heterogeneity of cell types in PSC, we analyzed cell type–specific marker gene expression to assess their spatial localization. In addition, cholangiocyte marker genes were used to approximate periductal regions. The distinct expression patterns of hepatocyte and cholangiocyte genes delineated the lobular and periductal regions of the liver (**Fig. S3, a and b**). Immune and stellate cell populations displayed distinct localization patterns; for example, LAM-like MP, MoMP and HSC1 were enriched in periductal regions, whereas HSC2 and KC were predominantly found in lobular regions occupied by hepatocytes (**Fig. S3, c-g**). In regions exhibiting onion-skin fibrosis, a characteristic feature of PSC (**Fig. 4a**), we observed concentric layers of LAM-like MP, MoMP, HSC1, and T cells surrounding the bile ducts (**Fig. 4b**). To quantitatively assess the differential spatial localization of these cell types, we performed a neighborhood enrichment analysis. We found that HSC1, LAM-like macrophages, MoMP, and CD4/8 T cells were significantly enriched around bile duct epithelial cells (at least 168% of expected, p-values < 0.0001), while HSC2 and KC were largely excluded from the cholangiocyte neighborhood (HSC2: 34% of expected; KC: 15% of expected, p-values < 0.0001, **Fig. 4c**). The clustered enrichment patterns suggested that PSC-associated cells formed two distinct niches. To further characterize these niches, we applied a cell niche discovery approach similar to that reported by CellCharter (15) and identified two spatially distinct cell niches (**Fig. 4d**). The first, termed the *fibrotic niche*, contained a large fraction of HSC1 cells and was enriched in LAM-like MP, MoMP, and CD4/8 T cells (**Fig. 4e**). This niche exhibited high expression of genes involved in collagen production, such as *COL3A1* and *COL1A1*, as well as *CCL21*, a chemoattractant for immune cells (**Fig. 4f**). In contrast, the second niche, termed the *non-fibrotic niche*, was predominantly composed of hepatocytes and displayed high expression of genes associated with normal liver metabolic functions, including *ADH1B* and *BAAT*. Notably, the spatial distribution of the fibrotic niche aligned with periductal regions.

**Figure 4.**
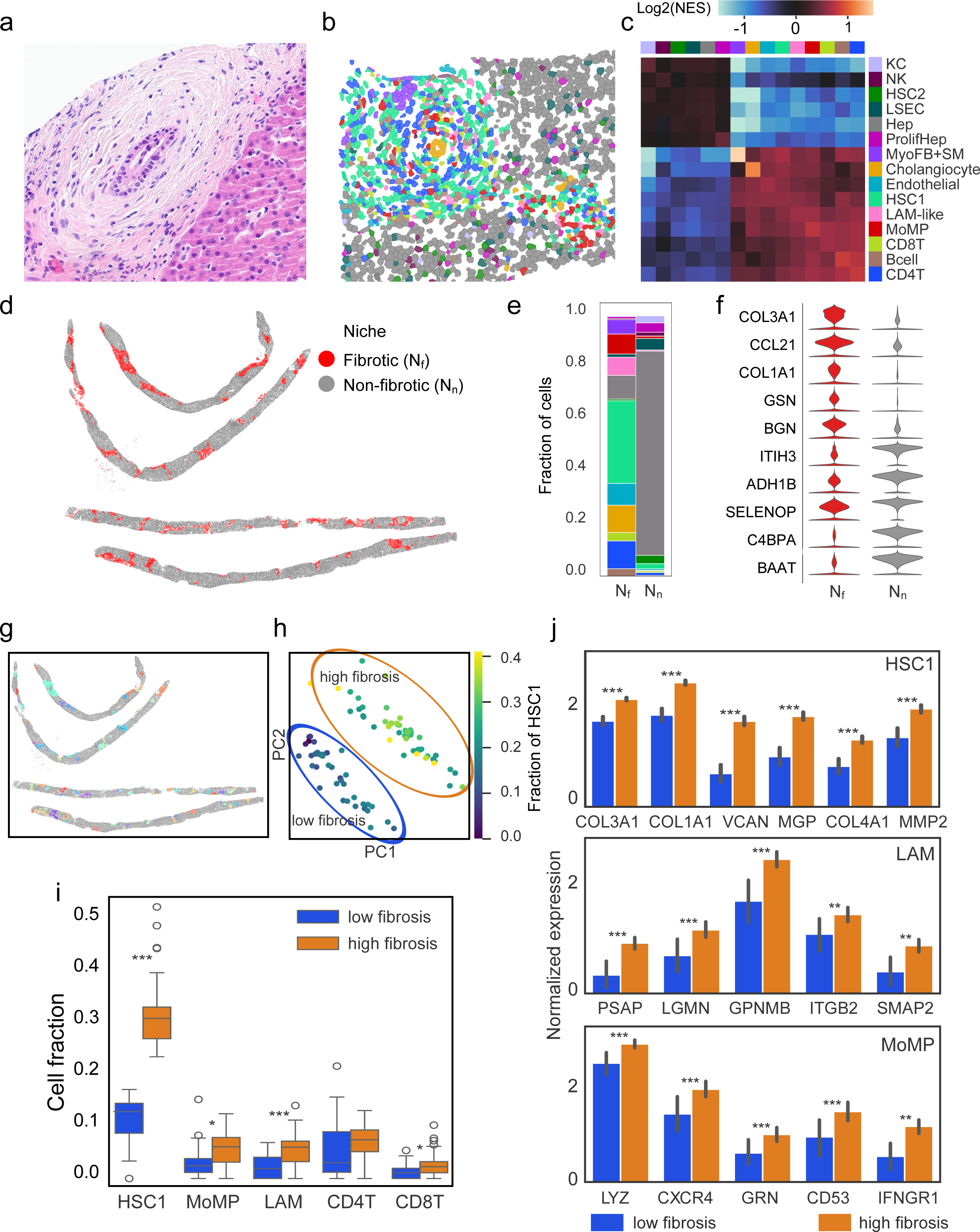
Neighborhood analysis of PSC fibrotic cell niches. (a) A region with onion skin fibrosis on a consecutive slice from the same tissue used in scSRT profiling; (b) A reconstructed image focusing on a region with onion skin fibrosis using scSRT data. Polygons in the image represent segmented cell masks and are colored based on cell types. (c) Heatmap showing neighborhood enrichment scores (NES) between each evaluated neighboring cell type (columns) in the proximity of the reference cell type (rows). Each cell in the heatmap is colored based on the log2(NES) value. (d) Reconstructed image showing the location of each cell in the scSRT data. Cells are shown as dots on the image and colored based on their niche identity. (e) Stacked bar plot showing the fraction of each cell type among all cells included the fibrotic and non-fibrotic niches. (f) Top 5 DEG for the fibrotic and non-fibrotic niches. Differential analysis was performed using the Wilcoxon’s rank sum test with FDR-adjusted p-value cutoff at 0.05. DEGs were ranked using log fold changes. (g) Reconstructed image of cells in scSRT data showing the location of periductal regions. (h) PCA plot calculated from the pseudobulk level expression profiles from periductal regions. Pseudobulks are colored based on its HSC1 fraction. (i) Boxplot showing the enrichment of cell types in the high-fibrosis regions compared to the low-fibrosis regions. Significance obtained from the student’s t test across the two groups. ∗p <0.05, ∗∗p <0.01, ∗∗∗p <0.001. (j) Stacked violin plot of genes upregulated in the high-fibrosis periductal regions. Differential analysis was performed in each cell type using the Wilcoxon’s rank sum test. DEG were filtered based on FDR-adjusted p-value cutoff at 0.05 and min. log fold change of 0.6.

### Periductal fibrosis correlates with enriched and transcriptionally reprogrammed HSC and macrophages

To better understand how the cellular microenvironment influences periductal fibrosis, we examined the cell type composition and gene expression within individual periductal regions within the fibrotic niche. We defined these regions by grouping cells within a 50-µm radius (equivalent to 4–5 surrounding cells) of a cholangiocyte into bins, excluding those that contained only isolated cholangiocytes. To prevent over-segmentation of a single periductal region, adjacent bins with overlapping cells were merged. This process identified 86 distinct periductal regions (**Fig. 4g**).

To assess the overall transcriptomic profiles of these regions, we pooled cells in each region into pseudobulks and performed unsupervised principal component analysis (PCA). This analysis revealed two distinct clusters, primarily driven by differences in the fraction of HSC1 cells (**Fig. 4h**). Based on this distinction, we categorized the regions as high- and low-fibrosis. Notably, HSC1 cells were 2.84-fold enriched in high-fibrosis regions, which also exhibited increased proportions of MoMP, LAM-like macrophages, and CD8 T cells (minimum fold enrichment = 1.74, **Fig. 4i**).

The enrichment of HSC1 cells in high-fibrosis periductal regions was accompanied by elevated expression of fibrosis-associated genes (*FDR-adjusted p* < 0.05, log fold change >= 0.6, **Fig 4j**). For example, *COL1A1, COL3A1,* and *COL4A1*—key markers of fibrosis known to be upregulated in PSC (16–18) showed significantly increased expression. Additionally, Versican *(VCAN)*, a component of the early fibroproliferative matrix that facilitates fibroblast migration and myofibroblast activation (19, 20), as well as macrophage activation via toll-like receptors (21), was upregulated. Notably, these genes exhibited an average 8-fold increase in HSC1 cells from high-fibrosis regions compared to those in low-fibrosis regions.

While macrophage populations in high- and low-fibrosis periductal regions showed no significant differences in the expression of genes linked to complement activation (*C1QA/B/C*) or the NF-κB activation complex (*TNFAIP2/3, MCL1, GPR183*), we observed distinct transcriptional changes associated with macrophage function (*FDR-adjusted p* < 0.05, log fold change >= 0.6). Specifically, genes involved in macrophage migration (*ITGB2, CXCR4, CD53*), activation (*IFNGR1, LYZ*), and fibrotic responses (*GPNMB, PSAP, LGMN*) were significantly upregulated in LAM-like and MoMP cells within high-fibrosis regions.

### CCC analysis predicts macrophages drive fibrogenic HSC activation

The close spatial association between HSC1 and macrophages, along with the upregulation of key fibrogenic genes in HSC1 within the fibrotic niche, suggests a functional link between the gene expression programs of these two cell types. To systematically investigate this relationship, we applied the Spacia algorithm to identify cell-cell communication (CCC) between HSC1 and its neighboring cells in the fibrotic niche (22). Spacia prioritizes single-cell-level CCCs by incorporating cell-cell proximity as a constraint and selecting interactions that lead to downstream changes in signal-receiving cells. We predicted CCCs by modeling HSC1 gene expression as a function of signal-sending genes expressed in its neighboring cells, including CD4/8 T cells, macrophages, and cholangiocytes. Genes selectively expressed in these neighboring cell types were considered candidate signal-sending genes, while genes upregulated in HSC1 served as response candidates. For each signal-response gene pair, we estimated two key parameters: the interaction score (β), which reflects the impact of the CCC on response gene expression, and the proximity coefficient (b), which indicates the dependency of CCC on cell proximity.

After filtering interactions based on statistical significance (p-values of β and b), we identified 236 HSC1-targeting CCCs originating from neighboring cells, with nearly half of these interactions driven by MoMP or LAM-like macrophages (**Fig. 5a**). Among the 20 genes in HSC1 upregulated by the sender cells, several were linked to fibrosis-related processes, including collagen formation (*COL1A1, COL1A2, COL3A1, COL6A1*), platelet-derived growth factor (PDGF) signaling (*PDGFRA, PDGFRB, THBS2*), extracellular matrix remodeling (*BGN, LUM, DPT, VCAN*), and chemotaxis (*CXCL12*). Interestingly, while T cells were associated with genes involved in extracellular organization and PDGF signaling, the major collagen genes linked to liver fibrosis (23)—*COL1A1, COL1A2,* and *COL3A1*—were predominantly associated with macrophage-derived signals (**Fig. 5b**).

**Figure 5.**
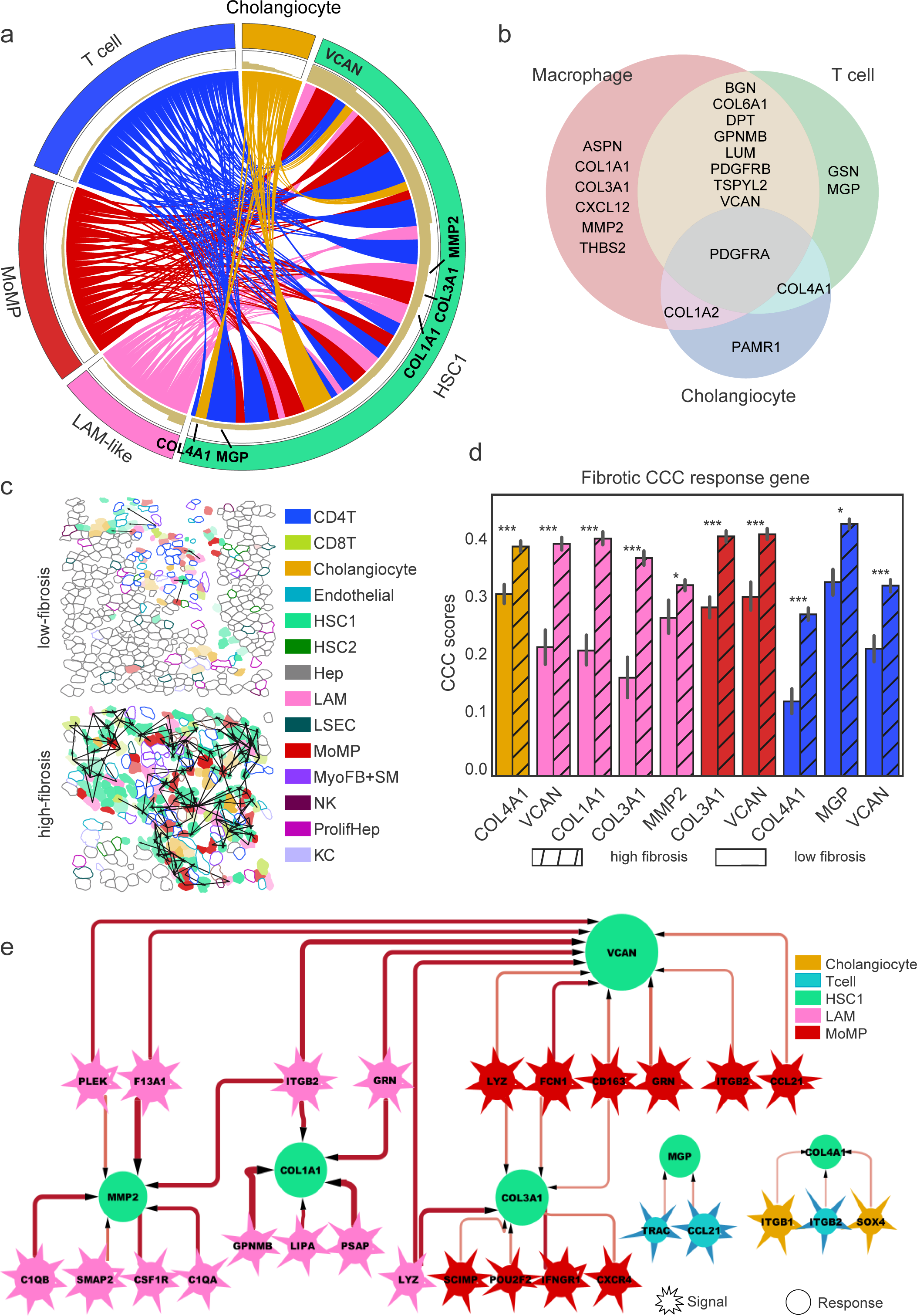
CCC analysis of periductal fibrotic niches in PSC. (a) Circos plot summarizing HSC1-targeting CCC from macrophages, T cells, and cholangiocytes. Each band represents a CCC between the sender cell and HSC1. The bar plot below each gene in the middle track represents the sum of absolute β values of each CCC involving this gene. CCC were predicted between HSC1 (receiver cell) and macrophages/T cells/cholangiocytes (sender cell). (b) Venn diagram showing the 20 HSC1 genes targeted by sender cells. (c) Reconstructed images of representative low- and high-fibrosis periductal regions. Polygons in the image represent segmented cell masks, and are colored based on cell types. CCC between cells are shown as directed arrows. (d) Comparison of primary instance (PI) scores in HSC1 fibrogenic genes between low- and high-fibrosis periductal regions. Standard deviations are shown as error bars. Colors of bars represent cell types shown in panel (c). ∗p <0.05, ∗∗p <0.01, ∗∗∗p <0.001. (e) CCC network focusing on HSC1 fibrogenic genes. Nodes with spiky borders represent genes expressed by sender cells, and nodes with smooth borders represent HSC1 genes. Nodes are colored based on cell types. CCC are shown as directed edges, and the absolute β values are positively proportional to edge thickness.

To assess the spatial distribution of these CCCs, we predicted the interaction probability between each macrophage, T cell, and cholangiocyte with HSC1, as estimated by the primary instance (PI) score in Spacia. As expected, these CCCs were restricted to cells in close proximity. Moreover, HSC1-targeting CCCs were confined to fibrotic niches in both liver samples and were enriched in high-fibrosis periductal regions (**Fig. 5c**), where HSC1 fibrotic genes—including COL1A1, COL3A1, COL4A1, MMP2, MGP, and VCAN—were upregulated. To further quantify the intensity of CCCs in high-versus low-fibrosis periductal regions, we compared the PI scores for each interaction between HSC1 and its neighboring cells. CCCs in high-fibrosis regions exhibited stronger interaction scores across all sender cell types, with macrophage-derived interactions targeting COL1A1, COL3A1, and VCAN displaying the most pronounced differences (**Fig. 5d**). A network representation of these interactions revealed that HSC1 fibrotic gene upregulation was predominantly driven by macrophages, with additional contributions from T cells and cholangiocytes (**Fig. 5e**). Specifically, COL1A1 was primarily activated by LAM-like macrophages and its expression in HSC1 was positively correlated with genes involved in macrophage activation (GPNMB), TLR signaling (ITGB2, doi:10.1002/eji.201242550; doi:10.1038/ni.1908), and fibroblast crosstalk (GRN, (24). Meanwhile, COL3A1 was mostly upregulated by MoMP and was positively associated with genes linked to inflammatory responses (IFNGR1, POU2F2) and cell recruitment (CXCR4). In summary, macrophage-HSC1 crosstalk is both enriched and intensified in high-fibrosis periductal regions and associated with upregulation of HSC1-derived fibrogenic genes.

### Orthogonal validation of fibrotic macrophages and HSC signatures

To validate the role of macrophages in driving fibrogenic gene expression in HSC1, we analyzed both bulk and scRNA-seq data from AILD patients, assessing the correlation between macrophage and HSC1 fibrotic gene programs and their association with PSC. Based on the predicted CCCs, we designed two gene signatures: one representing HSC1-derived fibrotic effector genes (COL1A1, COL3A1, COL4A1, MGP, MMP2, and VCAN) and another for macrophage-derived fibrotic driver genes (Table S3). To assess the activity of these gene signatures, we defined the fibrotic effector score as the average expression of the fibrotic effector genes, and the fibrotic driver score as the weighted average expression of the fibrotic driver genes, using Spacia-derived β coefficients as weights. We first calculated these scores in a public bulk RNA-seq dataset from adult AILD patients (25). In this dataset, both the fibrotic driver and effector scores were significantly higher in PSC patients compared to healthy controls (driver score P-value = 0.014; effector score P-value = 0.033, **Fig. 6a**). Similarly, in our in-house RNA-seq data from pediatric AILD patients, both scores were elevated in patients with advanced fibrosis (driver score P-value = 0.031; effector score P-value < 0.001, **Fig. 6b**).

**Figure 6.**
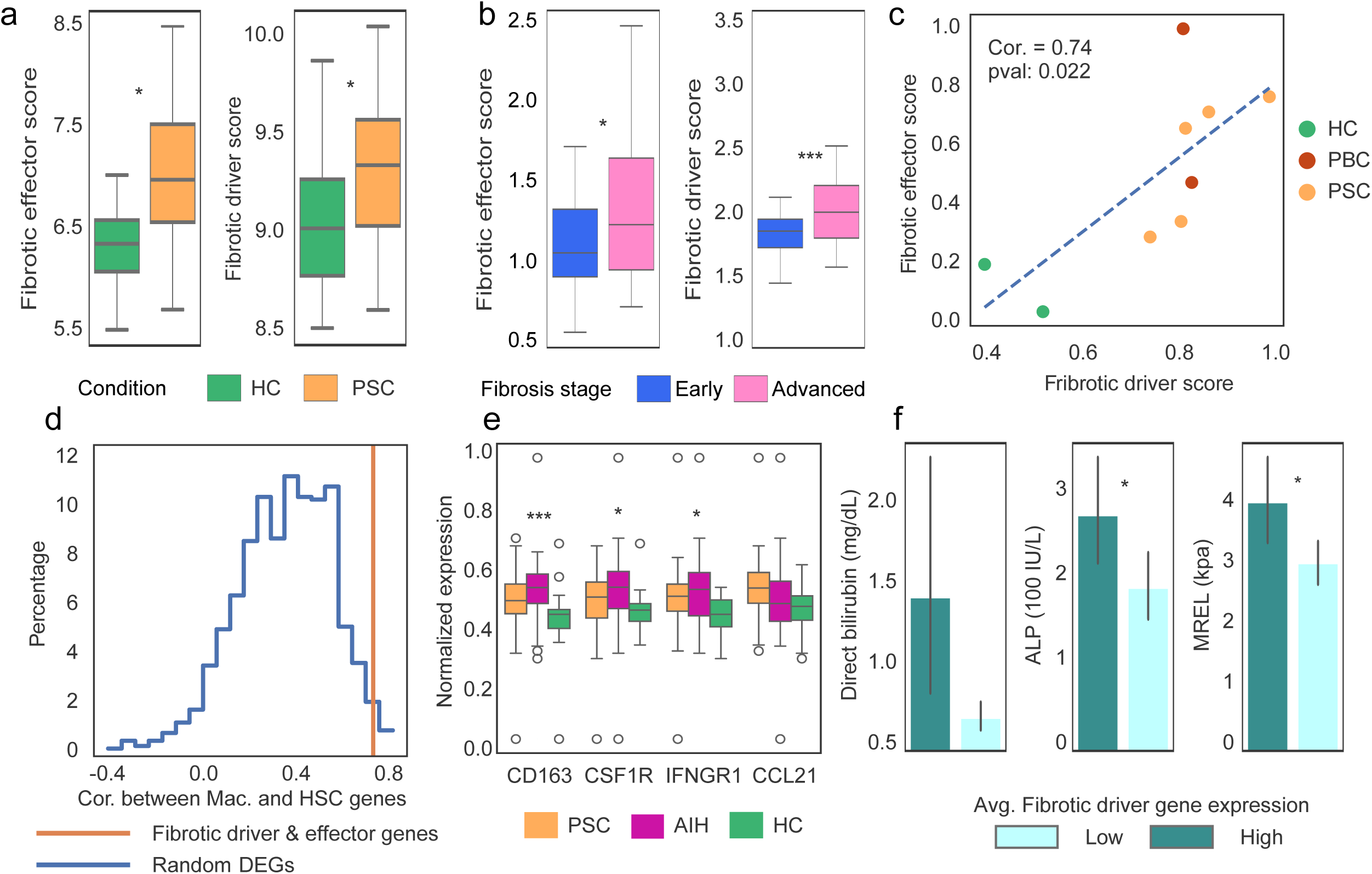
Orthogonal validation of effector and driver fibrotic signatures. (a) In silico validation of fibrotic driver and effector score in bulk liver RNA-seq data from healthy controls (HC) and adult patients with PSC. (b) Fibrotic driver and effector score in pediatric PSC patients with early or advanced stages of fibrosis, which is defined as METAVIR>=2 or Ishak >=3. (c) Pearson correlation between fibrotic driver scores calculated in macrophages and fibrotic effector scores calculated in fibroblasts. (d) Comparison between baseline correlation between macrophage- and fibroblast-DEGs, and correlation between the fibrotic driver and effector scores. Blue line represents the distribution of the baseline correlation in 1000 simulations using randomly sampled DEGs. Orange line represents the observed correlation between the fibrotic driver and effector scores, calculated in macrophage and fibroblasts, respectively. (e) Boxplot showing proteins that are elevated in PSC/AIH compared to HC in plasma. Statistical test was done using One-way Anova. (f) Bile duct injury and liver stiffness measures in AILD patients with high- or low-average plasma concentrations for fibrotic driver markers. All error bars represent standard deviation. ∗p <0.05, ∗∗p <0.01, ∗∗∗p <0.001.

Next, we leveraged a public scRNA-seq dataset from healthy and PSC/PBC donors (9) to examine the correlation between the fibrotic driver and effector gene signatures. The driver and effector scores were calculated at the single-cell level in macrophage and fibroblast populations, respectively, and then aggregated at the patient level. We observed a strong correlation between these scores (Pearson correlation = 0.74, P-value = 0.022, **Fig. 6c**). To ensure this correlation was not simply due to these genes being among the top expressed genes in macrophage and fibroblast populations, we conducted a control analysis using randomly sampled top DEGs specific to these cell types. While macrophage- and fibroblast-specific DEG expression was moderately correlated at the patient level (average Pearson correlation = 0.374), the observed correlation between the fibrotic driver and effector signatures was significantly stronger than this baseline (empirical P-value = 0.021, **Fig. 6d**). These findings suggest that the upregulation of macrophage fibrotic driver genes is specifically associated with increased expression of fibrotic genes in fibroblasts within the same patient.

To determine whether the macrophage fibrotic gene signature is also reflected at the protein level in plasma, we performed SomaScan-based proteomic profiling using plasma samples from healthy donors (HC, n=22), pediatric AIH- (n=43), and PSC patients (n=43; **Table 3**). 8 out of 17 macrophage fibrotic drivers (LYZ, GRN, GPNMB, FCN1, CD163, CSF1R, IFNGR1, and CCL21) were included in the SomaScan panel. Among them, plasma concentrations for CD163, CSF1R, IFNGR1, and CCL21 were increased in AILD patients compared to healthy controls (**Fig 6e**). Next, we calculated the fibrotic driver score based on the average expression of available proteins in the SomaScan panel. Patients were dichotomized based on this score, with those above the median assigned to the ‘high’ group and those below to the ‘low’ group. Clinical measures were compared between the two groups. Markers of hepatocellular injury, i.e., serum AST and ALT, did not differ between the groups (data not shown). However, biomarkers of bile duct epithelial injury—including serum direct bilirubin- and ALP-levels—were elevated in patients with higher fibrotic driver scores (direct bilirubin P-value = 0.052; ALP P-value = 0.025; **Fig 6f**).

Out of the 108 plasma samples, 63.5% were collected with a median of 0.35 months before or after a research MR elastography (MReL) examination. We had shown before that MReL-derived liver stiffness values predicted histological stage of fibrosis in pediatric AILD (26). In this study, patients with higher fibrotic driver scores exhibited significantly higher liver stiffness values by MReL (4.2 kPa vs. 3.1 kPa, P-value = 0.014).

## Discussion

Macrophages have been implicated as key drivers of liver fibrosis in adult end-stage liver disease. Building on these findings, we investigated macrophage-fibroblast crosstalk in pediatric-onset PSC across the disease severity spectrum. To this end, we applied an integrative multiomic approach—combining scSRT, bulk RNA-seq, and SomaScan plasma proteomics—to liver biopsies and plasma samples from children and adolescents with AILD. Using scSRT, we identified fibrotic periductal niches enriched in LAM-like MP and MoMP, but not Kupffer cells, alongside a population of highly activated HSC expressing high levels of fibrogenic genes. These niches also included cholangiocytes, T cells, and B cells, recapitulating classical PSC histopathology such as concentric fibrosis and ductular reaction. To account for spatial heterogeneity and to discern the cellular crosstalk between these cell types, we segmented periductal regions and applied Spacia to model CCC, predicting that macrophages expressing *LYZ, GRN, GPNMB, FCN1, CD163, CSF1R, IFNGR1*, and *CCL21* induce a profibrogenic transcriptional program in HSC, characterized by upregulation of *COL1A1*, *COL3A1, COL4A1, MGP, MMP2, and VCAN*. These macrophage-fibroblast interactions were validated across independent datasets, including adult PSC single-cell and bulk RNA-seq, and were strongly associated with advanced fibrosis in a pediatric AILD cohort (n=66). Furthermore, a plasma cytokine signature derived from these macrophage genes *(CD163, CSF1R, IFNGR1, CCL21*) correlated with elevated liver stiffness in 102 pediatric AILD patients, supporting their utility as non-invasive biomarkers. Collectively, our findings define a conserved profibrotic macrophage-fibroblast crosstalk in PSC and establish a robust framework for spatially resolved CCC analysis in fibro-inflammatory liver diseases.

MoMP has been linked to hepatocyte loss and ductal reaction (27). Notably, macrophage populations enriched around cholangiocytes in PSC livers have been reported to exhibit high C1QA/B, moderate CD163, and low-to-moderate TREM1 and TNF gene expression (28), aligning with the MoMP phenotype observed in the periductal regions of our scSRT data. This gene expression profile of MoMP suggests it has reduced inflammatory potential, which has been reported *in vitro* where PSC macrophages showed reduced capacity to produce TNF upon activation (9). LAM-like MP has been reported in PSC and cirrhosis (9, 10, 29), with high expression of genes including *CD9, TREM2, GPNMB and LGMN.* Consistent with our findings, these macrophages were shown to be enriched in periductal fibrotic regions in PSC livers (9). Given the enrichment of MoMP and LAM-like MP in HSC-rich periductal areas, we modeled intercellular crosstalk between these cell types. While spatial heterogeneity in gene expression often introduces analytical uncertainty, we leveraged this variability by focusing on HSC genes specifically upregulated in regions with elevated fibrosis. The cell-specific gene programs underlying this crosstalk were initially derived from a limited dataset (n=2). To address this limitation and enhance the robustness of our predictions, we validated these programs using two independent and larger patient cohorts, and further prioritized genes involved in macrophage-HSC interactions. Among the prioritized genes, CD163 is a well-established biomarker of poor prognosis in PSC (30, 31). CCL21 is upregulated in PSC and plays a role in the development of secondary lymphoid structures and chronic inflammation. It acts on CCR7, which is expressed in cultured HSC, promoting their activation *in vitro* (32); however, its profibrotic role in vivo has not been previously reported. Additionally, CSF1R has been proposed as a therapeutic target for inflammatory diseases (33), though its plasma levels have not been linked to liver fibrosis in PSC before.

scSRT is an emerging, promising tool to study pathogenesis mechanisms in disease, as it is robust against cell type biases imposed by the cell preparation steps that is common in single-cell RNA-seq, and allows highly desired analysis such as cell niche profiling and cell-proximity constrained CCC predictions. However, several data quality issues could limit this promise. The first limitation in using scSRT for CCC prediction is rooted from its dependency on cell segmentation. Errors in this process could lead to erroneous identification of co-expression within the same cell as CCC. In this study, we mitigate this issue by applying the Baysor segmentation algorithm and limiting the CCC search space to genes specifically expressed in sender/receiver cell types. Secondly, scSRT data is highly sparse and thus highly susceptible to the ’dropout’ effect typical in single-cell expression data. This issue can be mitigated by applying gene expression denoising or dropout correction tools to the scSRT data, such as Sprod, which we used in this study to correct gene expression noise.

Our results were derived and validated primarily using data from pediatric AILD patients, where co-occurrence of AIH in PSC is more common compared to adults. Additionally, the heterogeneity of PSC—such as the distinction between large duct and small duct PSC—poses challenges for interpretation. These factors may limit the generalizability of our findings in adult PSC patients. Like most studies involving scSRT, our discovery potential is limited to the predefined gene panels. However, with the emergence of whole transcriptome scSRT platforms such as CosMX WTX, future studies may expand our CCC findings by modeling the director effectors, such as cytokines, chemokines, and growth factors, and evaluate their impact on bile duct injury. Our results demonstrate the power of high-resolution scSRT to uncover novel pathogenic mechanisms through CCC modeling. We present a comprehensive analytical framework that addresses scSRT data quality challenges and incorporates cell niche analysis, refined segmentation, and robust CCC prediction. This framework offers a powerful and flexible strategy for dissecting cellular interactions in fibrotic disease and for advancing future spatial omics research.

## Methods

### Pediatric AILD Cohort

Patients with pediatric onset AILD receiving care at CCHMC were enrolled into the CALD observational study, registered as NCT03175471, between February of 2017 and July 2023. Following consent to the Institutional Review Board (IRB) approved study (IRB#2016-7388), blood was collected from patients with an established or suspected diagnosis of AIH or PSC. The clinical diagnosis of PSC was assigned based on established guidelines (2). Patients were assigned the diagnosis of AIH if they met the international autoimmune hepatitis study group (IAIHG) simplified criteria (34) and did not have radiologic or histological evidence of cholangiopathy. Research MRI examinations were performed on the AILD cohort (NCT03175471, IRB#2016-7388), as previously described (35). Briefly, MRI were performed at a field strength of 1.5 Tesla (Ingenia; Philips Healthcare, Best, the Netherlands). Axial 2D spin-echo echo-planar Magnetic Resonance Elastography (MRE) was performed through the mid liver at four levels and analyzed as previously described (35).

### VizGen MERSCOPE SRT data generation

Sample preparation. Needle biopsies from two PSC patients with early stage of disease were collected under IRB protocol IRB#2017-2284 and IRB#2016-7388. Freshly cut, unstained sections from archived FFPE liver biopsies of these two patients were mounted to slides provided by VizGen for hybridization according to the vendor’s instructions.

### SomaScan plasma proteomics profiling

108 plasma samples from 43 AIH, 43 PSC and 22 HC donors were used for the SomaScan (1.3k proteins assay) proteomic study. Data collection, protein quantification, and quality control steps were performed at the Genome Technology Access Center in the Department of Genetics at Washington University School of Medicine, as previously described (36, 37). SomaScan data was normalized using log and quantile transformation. MP fibrotic driver score was calculated by averaging the normalized expression of the involved proteins. SomaScan samples were then divided into two groups based on the median of the fibrotic driver scores. Mean testing of clinical measures was done using the student’s t test.

### RNA-seq data acquisition, processing and analysis

RNA-seq data acquisition. Excess liver tissue from clinically indicated liver biopsies from 34 PSC and 30 AIH patients were used for bulk liver RNA-seq under IRB IRB#2017-2284 and IRB#2016-7388. Liver tissue samples were stored in RNA later in -80C. RNA was isolated with the miRNeasy Mini Kit (Qiagen, Germany) according to the manufacturer’s instructions. RNA integrity was confirmed by Agilent assays. RNA-seq was performed by the University of Cincinnati DNA core using 101 base pair, paired end reads at a read depth of 50 million bp, as reported before (38)

### RNA-seq data preprocessing and analysis

The alignment of reads to the human reference genome (GRCh38) was done using STAR (39) (v2.7.2b). Transcript quantification was done using FeatureCounts (40) (v1.6.4). The R package DEseq2 (40) (v1.26) was used to normalize gene counts and to generate variance stabilized count matrix. The Python package scanpy (41) (v1.9) was used for unsupervised clustering and differential expression analysis. Cutoff values of absolute fold change greater than 1.5 and FDR<0.05 were used to select for differentially expressed genes between sample group comparisons. Gene Set Enrichment Analysis (GSEA) was performed using the Enrichr API (42) with the ‘Reactome_Pathways_2024’ reference library. Sets of genes representing macrophage subsets and activities (9) were added to this library. The reference scRNA-seq data set was download from CELLxGENE (https://cellxgene.cziscience.com/collections/0c8a364b-97b5-4cc8-a593-23c38c6f0ac5). UMAP embeddings were re-calculated using the 400 MERSCOPE genes.

### SRT Data preprocessing, normalization and quality control

#### Raw Image Data Preprocessing

Spatial gene expression was decoded using the MERlin pipeline (43) on the VizGen MERSCOPE platform (software version 234b). Image stacks from multiple MERFISH rounds were aligned, background noise was filtered, and RNA spot detection was enhanced. Individual RNA molecule barcodes were decoded using a pixel-based algorithm with an adaptive barcoding scheme that corrected misidentified barcodes not matching the provided codebook. Raw image tiles of DAPI (nuclear staining), poly-T (cytoplasmic and nuclear staining), and cell boundaries (cell membrane IHC) were compiled into mosaic images across 7 z-planes using MERSCOPE software.

#### Cell Segmentation

Cell segmentation was performed using a VizGen-optimized implementation of the Cellpose algorithm (44) on the DAPI and cell boundary images. For each segmented cell, a unique identifier, outline mask, x-y-z coordinates, and shape metrics were recorded. Transcript molecules were assigned to cells based on spatial overlap with the cell masks. The total number of transcripts per gene per cell was then aggregated into a cell-by-gene expression matrix for downstream analysis.

#### SRT Data preprocessing and quality control

Low quality cells identified as those with fewer than 20 transcripts were removed from further analysis. Library size normalization, unsupervised clustering, and differential expression analysis were performed using the Python scanpy package (41) (v1.9, 29409532). Baysor (11) (v0.7.1) was used to refine transcript-to-cell assignments.

Sprod was applied to log transformed, library size normalized data to reduce noise in scSRT data. Cell type annotation was performed using a reference PSC scRNA-seq dataset based on marker gene expressions (9).

### Neighborhood and niche analysis

#### Neighborhood enrichment analysis

The ratio between the observed neighboring cell type composition and the expected cell type composition estimated based on abundance of cell types were calculated as the enrichment score of cell type n in the neighborhood of cell type m. The expected fraction of neighboring cell types is estimated assuming cell types are uniformly distributed across the whole slide. Enrichment scores greater than 1 indicated enrichment and values less than 1 indicated exclusion. The significance of such enrichment was calculated using Fisher’s exact test. In this study, a cell’s neighboring cells were defined as those within 50um.

#### Cell niche discovery

In this study, a cell’s niche was defined as the collective gene expression profile of its surrounding neighborhood. For each cell, the niche was computed by averaging the PCA embeddings of all neighboring cells. This resulted in a cell-by-niche embedding matrix, which was subsequently clustered using the Leiden algorithm to identify groups of cells with similar composition and expression patterns in neighboring cells.

#### Periductal region segmentation

In this study, periductal regions were referred to as the cell neighborhood within 50um of clustered cholangiocytes, which represent bile ducts or ductal reaction. To avoid over segmentation, cholangiocyte neighborhoods sharing cells were merged into one periductal region. Characterization of each periductal region was done by evaluating the average gene expression of all cells in the region. These analyses were done with custom scripts using the following Python packages: scikit-learn, scipy, and shapely.

### CCC prediction

CCC was inferred using the Spacia algorithm, which models the association between the expression of a signal gene in sender cells and a response gene in receiver cells. For each receiver cell type of interest, spatial neighborhoods containing at least two sender cells of a specified type were identified, forming multiple multi-cell instances (referred to as “bags”). Each bag included one receiver cell and its surrounding sender cells. A Bayesian multi-instance learning (MIL) framework was then applied to estimate the probability and magnitude of CCC between the candidate signaling and response gene pairs within each bag. The receiver cell type was set to ‘HSC1’, and the sender cell type includes CD4/8 T cells, MoMP, LAM-like MP and cholangiocytes. Candidate response and signal genes were selected as genes differentially expressed in HSC1 and expressed in at least 20% of the cells. The radius of the cell neighborhood was set to 50um.

### Orthogonal validation

Four datasets were used in the *in silico* validation, including a public RNA-seq dataset from adult PSC patients and HC donors, an inhouse RNA-seq dataset from pediatric AILD patients, a sc/snRNA-seq dataset from PSC patients, and an inhouse SomaScan plasma proteomics dataset from pediatric AILD and HC donors. Normalized expression matrix of the adult RNA-seq dataset (25) was downloaded from the Gene Expression Omnibus (GEO: GSE61260). sc/snRNA-seq data of adult PSC patients was downloaded from CELLxGENE as previously mentioned. Count data was normalized using library size normalization and log transformation. In RNA-seq and SomaScan datasets, macrophage fibrogenic driver score was calculated as the average expression of the fibrotic driver genes, including *ITGB2, LYZ, GRN, F13A1, PLEK, GPNMB, FCN1, PSAP, CD163, C1QB, CXCR4, CSF1R, C1QA, LIPA, IFNGR1, SMAP2, and CCL21*.

HSC fibrogenic effector score was calculated similarly using the fibrotic effector genes, including COL1A1, COL3A1, COL4A1, MGP, MMP2, and VCAN. In the sc/snRNA-seq dataset, the macrophage fibrotic driver score was calculated as the weighted sum of the fibrogenic driver genes in macrophage populations, β values predicted by Spacia were used as the weights (Table S2).

## Statistics

For scRNA-seq and scSRT data, differential expression analysis was performed using the Wilcoxon rank sum test, with fold change cut off at 1.5 and p-value cut off at 0.05. Two-sided T-test was used to evaluate the statistical significance of macrophage signature expression between sample groups.

## Data Availability

The processed scSRT data including cell-by-gene counts, cell and transcript metadata are available at [**Accession number pending**]. Raw images associated with the study are available upon request. RNA-seq data from 64 AILD patients are available on GEO with accession number GSE303271.

## Author Contributions

Y.W. and A. G. M. conceived of the project, obtained the funding, performed the experiments, and wrote the manuscript. Y.W., D.A., X. X., and Z. Y. analyzed the RNA-seq, sc/snRNA-seq and scSRT data. C. CR and M.A. recruited the study participants, M. S., AyvH, L. P., M. A., and C. C. generated the RNA-seq and SomaScan data. P. S. performed pathological evaluations of patients involved in this study. J.R.D. and A.T.T. contributed to studying pathology images and reviewed the final manuscript.

## Funding Sources

This project was supported in part by National Institute of Health (NIH) R01DK095001 to Y.W and A.G.M, U01 DK62497 to A.G.M, the NIH P30 DK078392 (Bio-Imaging and Analysis Core, Single Cell Genomics Core, Integrated Pathology Research Core) of the Digestive Diseases Research Core Center in Cincinnati, the Center for Autoimmune Liver Disease (CALD) at CCHMC, and the Cincinnati Children’s Research Foundation through the 2024 Center for pediatric Genomics Award to Y.W.

**Figure S1. Top differentially expressed genes in PSC cell types.**

Dot plot illustrating the expression of the top 10 marker genes for each cell type. Dot size represents the proportion of cells expressing the gene, while color intensity indicates the expression level. Gene expression in log (CPM+1) was scaled to a range of 0 to 1.

**Figure S2. UMAP embeddings of PSC cell types from the reference dataset.** Recalculated UMAP embeddings of cells in the reference scRNA-seq (a) and snRNA-seq (b) dataset. Clusters are color-coded based on cell types.

**Figure S3: Spatial distribution of cell types identified in the scSRT PSC samples.** Panels (a–h) depict the spatial localization and average expression of cell top type-specific marker genes across FFPE liver tissue slides. Colors are mapped based on the average expression of the top 10 differentially expressed genes in each cell type. Cells are plotted based on the spatial coordinates of their centroids.

## Supporting information

All tables including supplementary tables

## REFERENCES

1. Liu JZ, Hov JR, Folseraas T, Ellinghaus E, Rushbrook SM, Doncheva NT, et al. Dense genotyping of immune-related disease regions identifies nine new risk loci for primary sclerosing cholangitis. Nat Genet. 2013;45(6):670–5.

2. Deneau MR, El-Matary W, Valentino PL, Abdou R, Alqoaer K, Amin M, et al. The natural history of primary sclerosing cholangitis in 781 children: A multicenter, international collaboration. Hepatology. 2017;66(2):518–27.

3. Laborda TJ, Jensen MK, Kavan M, and Deneau M. Treatment of primary sclerosing cholangitis in children. World J Hepatol. 2019;11(1):19–36.

4. Dotan M, Fried S, Har-Zahav A, Shamir R, Wells RG, and Waisbourd-Zinman O. Periductal bile acid exposure causes cholangiocyte injury and fibrosis. PLoS One. 2022;17(3):e0265418.

5. Ceci L, Gaudio E, and Kennedy L. Cellular Interactions and Crosstalk Facilitating Biliary Fibrosis in Cholestasis. Cell Mol Gastroenterol Hepatol. 2024;17(4):553–65.

6. Cadamuro M, Girardi N, Gores GJ, Strazzabosco M, and Fabris L. The Emerging Role of Macrophages in Chronic Cholangiopathies Featuring Biliary Fibrosis: An Attractive Therapeutic Target for Orphan Diseases. Front Med (Lausanne). 2020;7:115.

7. Guicciardi ME, Trussoni CE, Krishnan A, Bronk SF, Lorenzo Pisarello MJ, O’Hara SP, et al. Macrophages contribute to the pathogenesis of sclerosing cholangitis in mice. J Hepatol. 2018;69(3):676–86.

8. De Muynck K, Vanderborght B, De Ponti FF, Gijbels E, Van Welden S, Guilliams M, et al. Kupffer Cells Contested as Early Drivers in the Pathogenesis of Primary Sclerosing Cholangitis. Am J Pathol. 2023;193(4):366–79.

9. Andrews TS, Nakib D, Perciani CT, Ma XZ, Liu L, Winter E, et al. Single-cell, single-nucleus, and spatial transcriptomics characterization of the immunological landscape in the healthy and PSC human liver. J Hepatol. 2024;80(5):730–43.

10. Ramachandran P, Dobie R, Wilson-Kanamori JR, Dora EF, Henderson BEP, Luu NT, et al. Resolving the fibrotic niche of human liver cirrhosis at single-cell level. Nature. 2019;575(7783):512–8.

11. Petukhov V, Xu RJ, Soldatov RA, Cadinu P, Khodosevich K, Moffitt JR, et al. Cell segmentation in imaging-based spatial transcriptomics. Nat Biotechnol. 2022;40(3):345–54.

12. Poch T, Krause J, Casar C, Liwinski T, Glau L, Kaufmann M, et al. Single-cell atlas of hepatic T cells reveals expansion of liver-resident naive-like CD4(+) T cells in primary sclerosing cholangitis. J Hepatol. 2021;75(2):414–23.

13. Arceneaux D, Chen Z, Simmons AJ, Heiser CN, Southard-Smith AN, Brenan MJ, et al. A contamination focused approach for optimizing the single-cell RNA-seq experiment. iScience. 2023;26(7):107242.

14. Denisenko E, Guo BB, Jones M, Hou R, de Kock L, Lassmann T, et al. Systematic assessment of tissue dissociation and storage biases in single-cell and single-nucleus RNA-seq workflows. Genome Biol. 2020;21(1):130.

15. Varrone M, Tavernari D, Santamaria-Martínez A, Walsh LA, and Ciriello G. CellCharter reveals spatial cell niches associated with tissue remodeling and cell plasticity. Nat Genet. 2024;56(1):74–84.

16. Saffioti F, Hall A, de Krijger M, Verheij J, Hübscher SG, Maurice J, et al. Collagen proportionate area correlates with histological stage and predicts clinical events in primary sclerosing cholangitis. Liver Int. 2021;41(11):2681–92.

17. Vesterhus M, Nielsen MJ, Hov JR, Saffioti F, Manon-Jensen T, Leeming DJ, et al. Comprehensive assessment of ECM turnover using serum biomarkers establishes PBC as a high-turnover autoimmune liver disease. JHEP Rep. 2021;3(1):100178.

18. Thorburn D, Leeming DJ, Barchuk WT, Wang Y, Lu X, Malkov VA, et al. Serologic extracellular matrix remodeling markers are related to fibrosis stage and prognosis in a phase 2b trial of simtuzumab in patients with primary sclerosing cholangitis. Hepatol Commun. 2024;8(7).

19. Bensadoun ES, Burke AK, Hogg JC, and Roberts CR. Proteoglycan deposition in pulmonary fibrosis. Am J Respir Crit Care Med. 1996;154(6 Pt 1):1819–28.

20. Hattori N, Carrino DA, Lauer ME, Vasanji A, Wylie JD, Nelson CM, et al. Pericellular versican regulates the fibroblast-myofibroblast transition: a role for ADAMTS5 protease-mediated proteolysis. J Biol Chem. 2011;286(39):34298–310.

21. McLaren CE, Wagstaff M, Brittenham GM, and Jacobs A. Detection of two-component mixtures of lognormal distributions in grouped, doubly truncated data: analysis of red blood cell volume distributions. Biometrics. 1991;47(2):607–22.

22. Zhu J, Wang Y, Chang WY, Malewska A, Napolitano F, Gahan JC, et al. Mapping cellular interactions from spatially resolved transcriptomics data. Nat Methods. 2024;21(10):1830–42.

23. Kershenobich Stalnikowitz D, and Weissbrod AB. Liver fibrosis and inflammation. A review. Ann Hepatol. 2003;2(4):159–63.

24. Guerra RR, Kriazhev L, Hernandez-Blazquez FJ, and Bateman A. Progranulin is a stress-response factor in fibroblasts subjected to hypoxia and acidosis. Growth Factors. 2007;25(4):280–5.

25. Horvath S, Erhart W, Brosch M, Ammerpohl O, von Schönfels W, Ahrens M, et al. Obesity accelerates epigenetic aging of human liver. Proc Natl Acad Sci U S A. 2014;111(43):15538–43.

26. Dillman JR, Trout AT, Taylor AE, Khendek L, Kasten JL, Sheridan RM, et al. Association Between MR Elastography Liver Stiffness and Histologic Liver Fibrosis in Children and Young Adults With Autoimmune Liver Disease. AJR Am J Roentgenol. 2024;223(1):e2431108.

27. Guillot A, Winkler M, Silva Afonso M, Aggarwal A, Lopez D, Berger H, et al. Mapping the hepatic immune landscape identifies monocytic macrophages as key drivers of steatohepatitis and cholangiopathy progression. Hepatology. 2023;78(1):150–66.

28. Stankey CT, Bourges C, Haag LM, Turner-Stokes T, Piedade AP, Palmer-Jones C, et al. A disease-associated gene desert directs macrophage inflammation through ETS2. Nature. 2024;630(8016):447–56.

29. Miyamoto Y, Kikuta J, Matsui T, Hasegawa T, Fujii K, Okuzaki D, et al. Periportal macrophages protect against commensal-driven liver inflammation. Nature. 2024;629(8013):901–9.

30. Bossen L, Vesterhus M, Hov JR, Färkkilä M, Rosenberg WM, Møller HJ, et al. Circulating Macrophage Activation Markers Predict Transplant-Free Survival in Patients With Primary Sclerosing Cholangitis. Clin Transl Gastroenterol. 2021;12(3):e00315.

31. Elger T, Fererberger T, Huss M, Sommersberger S, Mester P, Stoeckert P, et al. Urinary soluble CD163 is a putative non-invasive biomarker for primary sclerosing cholangitis. Exp Mol Pathol. 2024;137:104900.

32. Bonacchi A, Petrai I, Defranco RM, Lazzeri E, Annunziato F, Efsen E, et al. The chemokine CCL21 modulates lymphocyte recruitment and fibrosis in chronic hepatitis C. Gastroenterology. 2003;125(4):1060–76.

33. Xiang C, Li H, and Tang W. Targeting CSF-1R represents an effective strategy in modulating inflammatory diseases. Pharmacol Res. 2023;187:106566.

34. Kaya M, Angulo P, and Lindor KD. Overlap of autoimmune hepatitis and primary sclerosing cholangitis: an evaluation of a modified scoring system. J Hepatol. 2000;33(4):537–42.

35. Dillman JR, Serai SD, Trout AT, Singh R, Tkach JA, Taylor AE, et al. Diagnostic performance of quantitative magnetic resonance imaging biomarkers for predicting portal hypertension in children and young adults with autoimmune liver disease. Pediatr Radiol. 2019;49(3):332–41.

36. Lertudomphonwanit C, Mourya R, Fei L, Zhang Y, Gutta S, Yang L, et al. Large-scale proteomics identifies MMP-7 as a sentinel of epithelial injury and of biliary atresia. Sci Transl Med. 2017;9(417).

37. Candia J, Cheung F, Kotliarov Y, Fantoni G, Sellers B, Griesman T, et al. Assessment of Variability in the SOMAscan Assay. Sci Rep. 2017;7(1):14248.

38. Lam S, Singh R, Dillman JR, Trout AT, Serai SD, Sharma D, et al. Serum Matrix Metalloproteinase 7 Is a Diagnostic Biomarker of Biliary Injury and Fibrosis in Pediatric Autoimmune Liver Disease. Hepatol Commun. 2020;4(11):1680–93.

39. Dobin A, Davis CA, Schlesinger F, Drenkow J, Zaleski C, Jha S, et al. STAR: ultrafast universal RNA-seq aligner. Bioinformatics. 2013;29(1):15–21.

40. Love MI, Huber W, and Anders S. Moderated estimation of fold change and dispersion for RNA-seq data with DESeq2. Genome Biol. 2014;15(12):550.

41. Wolf FA, Angerer P, and Theis FJ. SCANPY: large-scale single-cell gene expression data analysis. Genome Biol. 2018;19(1):15.

42. Kuleshov MV, Jones MR, Rouillard AD, Fernandez NF, Duan Q, Wang Z, et al. Enrichr: a comprehensive gene set enrichment analysis web server 2016 update. Nucleic Acids Res. 2016;44(W1):W90–7.

43. Allen WE, Blosser TR, Sullivan ZA, Dulac C, and Zhuang X. Molecular and spatial signatures of mouse brain aging at single-cell resolution. Cell. 2023;186(1):194–208.e18.

44. Stringer C, Wang T, Michaelos M, and Pachitariu M. Cellpose: a generalist algorithm for cellular segmentation. Nat Methods. 2021;18(1):100–6.

