## Supplementary figures and images for "Macrophages drive a fibrogenic gene program of periductal fibroblasts in pediatric primary sclerosing cholangitis"

### Supplement Figure 1

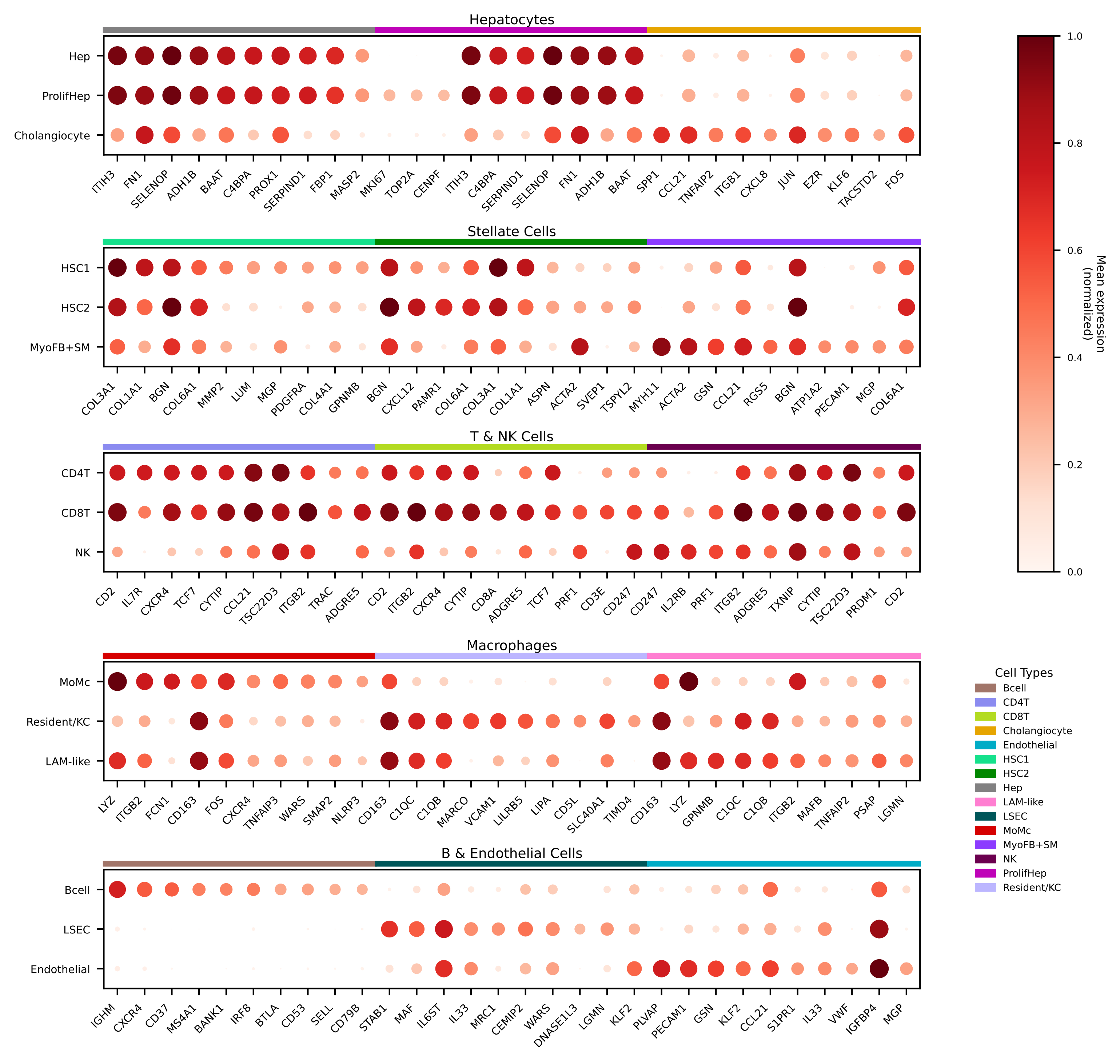

### Supplement Figure 2

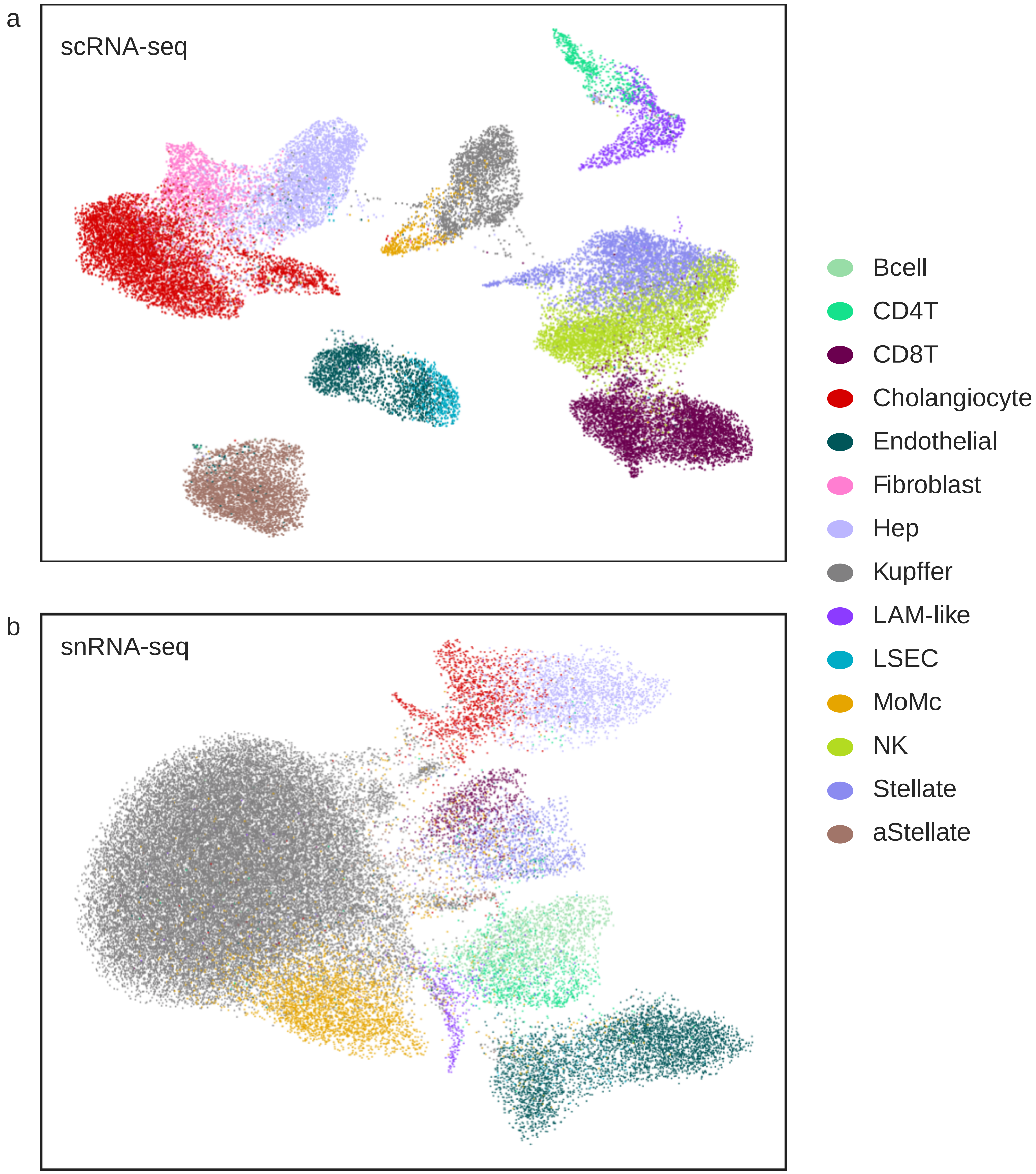

### Supplement Figure 3

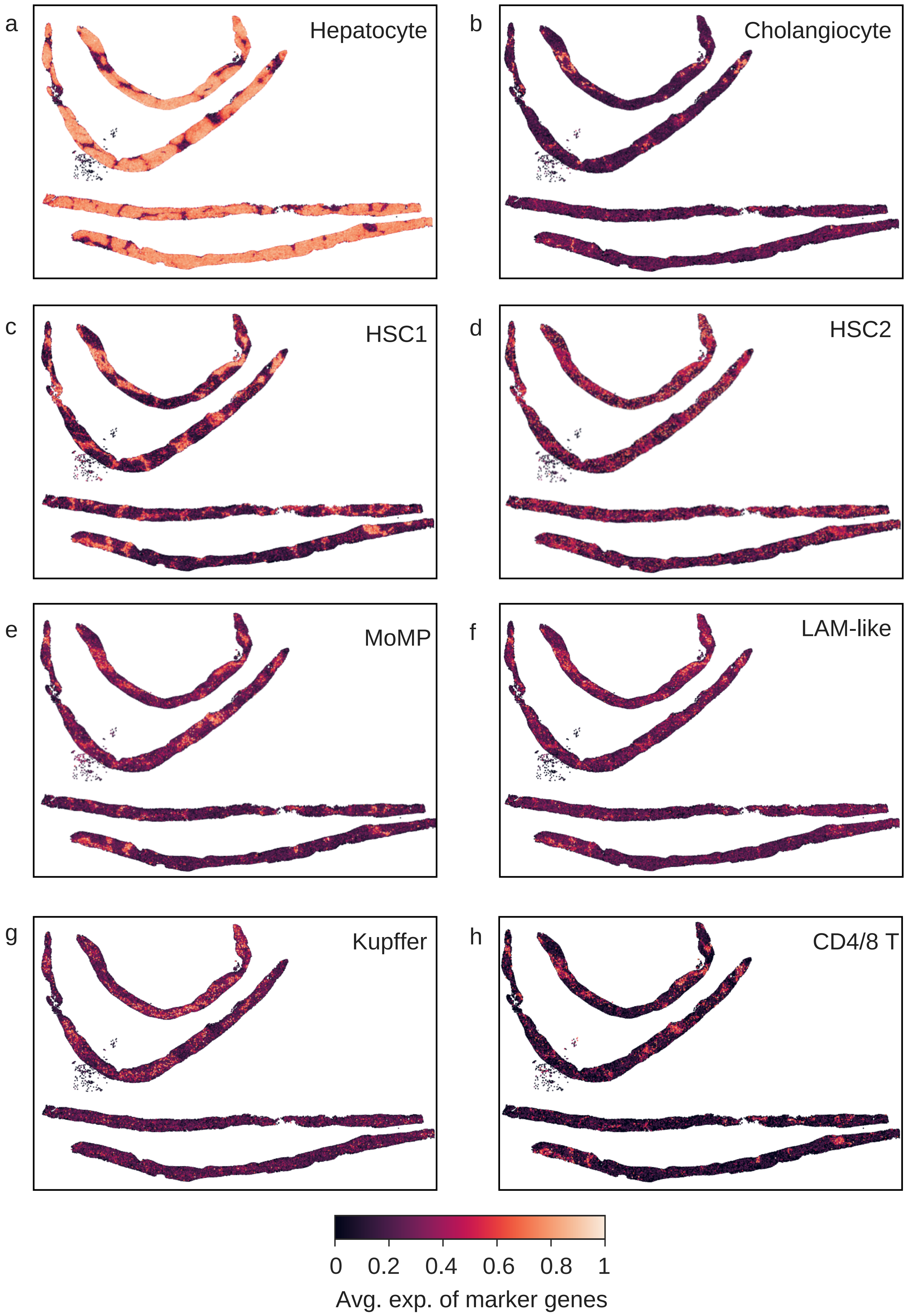
